# Spring temperature and land use change are associated with *Rana temporaria* reproductive success and phenology

**DOI:** 10.1101/2024.01.12.575320

**Authors:** Kat E. Oliver, Xavier A. Harrison

## Abstract

Chemical pollution, land cover change, and climate change have all been established as important drivers of amphibian reproductive success and phenology. However, little is known about the relative impacts of these anthropogenic stressors, nor how they may interact to alter amphibian population dynamics. Addressing this gap in our knowledge is important, as it allows us to identify and prioritise the most needed conservation actions. Here, we use long-term datasets to investigate landscape-scale drivers of variation in the reproductive success and phenology of UK Common frog (*Rana temporaria*) populations. Consistent with predictions, we found that increasing mean temperatures resulted in clear advancements in amphibian breeding phenology: earlier congregation of breeding *R. temporaria*, earlier initialisation of spawning, and earlier hatching. Temperature and number of frost days also affected rates of spawn mortality. However, temperature increases were also strongly correlated with increases in urban area, arable area, and nitrate levels in the vicinity of spawning grounds. None of these variables could explain variation in the total surface area of spawn present at breeding sites. These findings support previous work linking warming temperatures to shift in amphibian breeding phenology, but also highlight the importance of assessing the effect of land use change and pollution on wild amphibian populations. These results have implications for our understanding of the response of wild amphibian populations to climate change, and the management of human-dominated landscapes for declining wildlife populations.

## INTRODUCTION

Anthropogenic stressors are altering ecosystems across the world (Ceballos *et al*. 2015), with chemical pollution, land cover change, and climate change being highlighted as significant drivers of declines in ecosystem functioning (Wilcove *et al*. 1998; Walther *et al*. 2002; Howard *et al*. 2020). These altered ecosystems arise due to the effects of anthropogenic stressors on individual organisms, which can change community structures through decreased reproductive success and altered phenology (Nagelkerken & Munday 2016). Although anthropogenic stressors influence a myriad of organisms, amphibians are particularly susceptible due to their unique life cycle and physiology (Blaustein & Kiesecker 2002). Amphibia are subject to multiple threats including pathogens, land use change and climate change (Hof et al 2011). The combined effects of these processes have led to amphibians becoming the most threatened vertebrate group (Hoffmann *et al*. 2010), and their phenologies advancing at a fast rate compared to other vertebrate taxa (Parmesan 2006; Cohen *et al*. 2018). It is vital that we understand the drivers of these changes, and how these drivers may interact, as amphibians play many key roles in ecosystems (Burton & Likens 1975; Davic & Welsh Jr 2004; Mallory & Richardson 2005; Wood & Richardson 2010).

Climate change has affected amphibian reproductive success and phenology in both positive and negative ways. Increased winter precipitation resulting from climate change can benefit some amphibians (Benard 2015), but extreme weather events and advanced phenologies are likely to negatively impact others (Blaustein *et al*. 2010; Buss et al 2021). In a review of the impacts of increased droughts and extreme precipitation, Walls *et al*. (2013) conclude that even amphibian species adapted to variable environmental conditions are not able to adapt fast enough to keep up with dramatic changes in precipitation patterns. Drought negatively affects the reproductive success of amphibians, whilst warming temperatures are responsible for trends towards earlier amphibian breeding (Ficetola & Maiorano 2016). For example, warmer winter temperatures caused earlier breeding in *Rana sylva1ca*, but also lower female fecundity (Benard 2015), and have been associated with decreases in female body condition in female *Bufo bufo* (Reading 2007).

Chemical pollutants can increase mortality and alter rates of development in amphibian species (Carey & Bryant 1995), and the release of many chemical pollutants into the environment is growing (Sharma *et al*. 2020). The application of nitrate and ammonium fertilisers in the United States has increased by 4000% since the 1940s (Cao *et al*. 2018), which has led to environmental nitrogen levels being high enough to impact amphibians (Rouse *et al*. 1999). Nitrogen fertilisers are toxic to amphibian embryos and larvae above certain levels, causing methemoglobinemia (Huey & Beitinger 1980), and can also lead to eutrophication (Boyer & Grue 1995). These effects can scale up to change reproductive success and phenology in amphibian populations. Ammonium nitrate exposure has been linked to species-specific survival in three Australian amphibian species (Hamer *et al*. 2004). Exposure of *Litoria aurea* larvae to 10-15 mg/l ammonium nitrate resulted in significantly reduced survival, but no such effect was seen in *Crinia signifera* or *Limnodynastes peronii* (Hamer *et al*. 2004). This suggests that the declines solely seen in *L. aurea* populations were due to the effect of ammonium nitrate on this species alone. Ammonium nitrate may also be responsible for declines in European species; *Hyla arborea*, *Discoglossus galganoi*, and *Bufo bufo*, all of which have reduced survival in <200 mg/l of ammonium nitrate (Ortiz *et al*. 2004). Ammonium nitrate lowers larval development rate of *Pleurodeles waltl*, *Bufo calamita*, and *Pelobates cultripes*, suggesting that sub-lethal doses of ammonium and nitrate ions could alter amphibian phenology by delaying metamorphosis (Ortiz *et al*. 2004).

The most likely driver of increased exposure to chemicals like ammonium nitrate is land cover change. Many amphibian species use terrestrial habitats to disperse from natal ponds (Semlitsch 2008), meaning high-quality terrestrial habitat is needed to prevent isolation (Marsh & Trenham 2001). Isolation can lead to reduced reproductive success in amphibian populations (Allentoft & O’Brien 2010), so both habitat quality and connectivity are vital, but land cover change can alter both these factors. Previous studies have shown that urbanisation reduces the area of suitable habitat available to many amphibians (Price *et al*. 2012), as well as significantly increasing fragmentation and isolation (Natuhara & Zheng 2022). Expansion of intensive agriculture can also lead to amphibian declines, with areas of Mediterranean cropland having significantly lowered amphibian abundance (Beja & Alcazar 2003) compared to the surrounding countryside. The mechanisms driving these changes are likely the combined effects of reduced connectivity of habitat fragments, alongside increased exposure to chemicals.

Despite this abundance of research on the impacts of individual anthropogenic stressors on amphibian reproductive success and phenology, few studies have evaluated the relative impacts of these stressors simultaneously. This deficit can lead to difficulties in identifying the most effective actions needed to conserve amphibian species and the ecosystems they belong to (Frick *et al*. 2020). Here we investigate the relative impacts of chemical pollutants, land cover change, and climate change on the reproductive success and phenology of one amphibian species: *Rana temporaria*.

*R. temporaria* is an Anuran species widely distributed across Europe (Dabagyan & Sleptsova 1991; Sillero *et al*. 2014) that breeds in still, fresh water (Haapanen 1982) but spend the majority of time in terrestrial habitats (Dabagyan & Sleptsova 1991). Although widespread, many *R. temporaria* populations are declining (Cooke 1972; Loman & Andersson 2007; Guarino *et al*. 2008) and their breeding phenology is advancing (Scott *et al*. 2008). Like other amphibian species, it seems that chemical pollutants, land cover change and climate change are altering *R. temporaria* reproductive success and phenology.

Although there are conflicting results as to whether ecologically relevant levels of ammonium and nitrate are directly lethal to *R. temporaria* spawn and larvae (Oldham *et al*. 1997; Johansson *et al*. 2001), multiple studies demonstrate that high levels of these ions can reduce larval *R. temporaria* fitness, as well as delay metamorphosis (Johansson *et al*. 2001; Manson 2002; Oromí *et al*. 2009). In contrast, land cover change has been shown to cause *R. temporaria* population declines due to reduced reproductive success. For example, Cooke (1972) attributes the national declines in UK *R. temporaria* populations between 1940 and 1970 to the draining of wetlands to make way for intensive agriculture and urbanisation. Similar impacts of agricultural expansion have been seen in Sweden, where cropland populations are declining (Loman & Andersson 2007). These declines are not due to increased chemical pollutants (Loman & Lardner 2006) but could be due to lack of the suitable terrestrial refugia that adults need (Marnell 1998), leading the reduced gene pools and reproductive success (Allentoft & O’Brien 2010). There is little evidence to suggest that climate change is causing similar declines, with the only example being a period of high temperatures leading to reduced female fecundity in France (Neveu 2009). On the other hand, warming temperatures have been widely associated with earlier breeding phenology in *R. temporaria* (Neveu 2009), with Scott *et al*. (2008) finding that dates of breeding congregations, spawning, and hatching advanced in correlation with warming temperatures.

Here, we use a 21-year long-term monitoring dataset of *R. temporaria* breeding populations in the UK to quantify the relative importance of chemical pollutants, land cover change, and climate change on *R. temporaria* reproductive success and phenology. We test the predictions that:

1. Increasing average temperatures associated with climate change will drive earlier breeding phenology, including congregation to breed, spawning and hatching.
2. Land cover change, measured as the expansion of arable and urban areas, will be associated with decreased reproductive success.
3. Increased levels of pollutants such as ammonium and nitrate will be associated with greater degrees of spawn mortality

## METHODS

### Data Sources

We obtained data on *R. temporaria* reproductive success and phenology, as well as ammonium and nitrate ion concentrations, from the Environmental Change Network (ECN) (Rennie 2017). Across the UK, there are 10 ECN sites which collect this data (Figure 1a). The ECN sites differed in the number of ponds sampled and the duration they were sampled for (Table 1).

**Figure 1.**
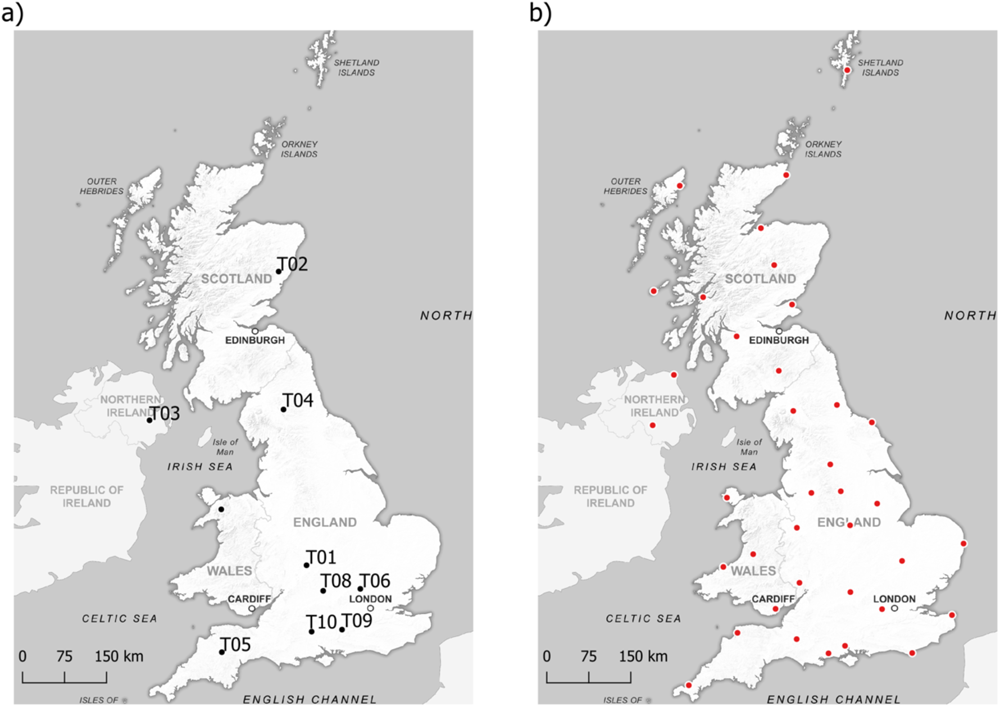
a) Locations of ponds sampled by the Environmental Change Network to obtain data on *R. temporaria* reproductive success and phenology, and agricultural pollutants, b) Locations of weather stations the Met Office use to measure climatic variables

**Table 1:** The number of ponds per ECN site and the years each ECN site was sampled for.

| ECN site | Co-ordinates | Number of ponds | First sampling year | Last sampling year |
| --- | --- | --- | --- | --- |
| T01 | 52.19361, -1.76417 | 1 | 1994 | 2012 |
| T02 | 56.90917, -2.55333 | 2 | 1994 | 2014 |
| T03 | 54.45333, -6.07806 | 2 | 1994 | 2011 |
| T04 | 54.695, -2.38778 | 7 | 1994 | 2015 |
| T05 | 50.78167, -3.91778 | 1 | 1994 | 2015 |
| T06 | 51.80333, -0.3725 | 1 | 1994 | 2010 |
| T08 | 51.781111, -1.33583 | 1 | 1994 | 1994 |
| T09 | 51.15444, -0.86306 | 1 | 1995 | 1995 |
| T10 | 51.12694, -1.63972 | 1 | 2001 | 2001 |
| T11 | 53.07455, -4.03351 | 2 | 1995 | 1995 |

From 1^st^ January every year, ponds were sampled weekly until males first congregated at the pond. This date was recorded, and subsequent sampling frequencies increased to daily until spawn first hatched. The dates of spawning and hatching were recorded. After hatching, sampling frequencies decreased to weekly until 16 weeks after spawning or when froglets were seen leaving the ponds. The date of leaving was recorded. The areas of spawn present and the percentage of spawn found dead were recorded when each pond was sampled. The concentrations of ammonium and nitrate ions were measured from the date of spawning by taking 250ml of pond water to analyse in a laboratory when each pond was sampled. We obtained data on land cover change from the UK Centre for Ecology & Hydrology (UKCEH) (Rowland 2020a, b). Using satellite data, the dominant land cover type in each 25x25m square of the UK in 1990 and 2015 were classified into 6 classes: woodland, arable, grassland, water, built-up areas, and other.

We obtained data on climate change from 37 historic Met Office stations across the UK (Figure 1b), which have been collecting climate data for at least 44 years (Met Office 2022). Climate data consists of monthly mean daily maximum temperature (°C), mean daily minimum temperature (°C), days with air frost, and total rainfall (mm).

### Data Processing

To quantify the land cover change surrounding each ECN location and the distances between each ECN location and each Met Office station, we used QGIS (QGIS Development Team 2022). Coordinates for the ECN locations were provided by the ECN, and the coordinates for the Met Office stations were taken from the Historic station data webpage (Met Office 2022), using the coordinate reference system OSGB 1996/British National Grid. To calculate the land cover change surrounding each ECN location we created a buffer zone with a radius of 2.5km surrounding each ECN location, as some common frogs can disperse over 2km (Kovar *et al*. 2009). We then used the land cover data obtained from UKCEH to calculate the area of each land cover class within these buffers in both 1990 and 2015. To calculate the distances between each ECN location and each Met Office station, we used the Distance matrix tool.

We used the software *R* (R Core Team 2020) to create a dataframe for each reproductive success and phenological response variable measured by the ECN (Table 2). Here, each row represented a pond at an ECN location in a single year from 1994-2015. The reproductive success data frames contained the largest surface area of spawn and percentage of dead spawn at each pond in each year. The phenology data frames contained the earliest date of either congregation, spawning, hatching, or leaving at each pond in each year, all converted to Julian days.

**Table 2:** The variables extracted for analysis and their sources. Abbreviations are used in the Principal Components Analysis plots.

| Extracted variable | Abbreviation | Unit of measurement | Data source |
| --- | --- | --- | --- |
| Congregation date | cong_date | Julian days | Rennie *et al.*, 2017 |
| Spawning date | spawn_date | Julian days | Rennie *et al.*, 2017 |
| Hatching date | hatch_date | Julian days | Rennie *et al.*, 2017 |
| Leaving date | leave_date | Julian days | Rennie *et al.*, 2017 |
| Surface area of spawn | surf_area | m<sup>2</sup> | Rennie *et al.*, 2017 |
| Percentage of spawn dead | perc_dead | % | Rennie *et al.*, 2017 |
| Ammonium concentration | spawn_nh4n | mg/l | Rennie *et al.*, 2017 |
| Nitrate concentration | spawn_no3n | mg/l | Rennie *et al.*, 2017 |
| Woodland area | woodland_area | m<sup>2</sup> | Rowland, 2020a<br>Rowland, 2020b |
| Arable area | arable_area | m<sup>2</sup> | Rowland, 2020a<br>Rowland, 2020b |
| Grassland area | grassland_area | m<sup>2</sup> | Rowland, 2020a<br>Rowland, 2020b |
| Freshwater body area | freshwater_area | m<sup>2</sup> | Rowland, 2020a<br>Rowland, 2020b |
| Built-up area | urban_area | m<sup>2</sup> | Rowland, 2020a<br>Rowland, 2020b |
| Other land cover area | other_area | m<sup>2</sup> | Rowland, 2020a<br>Rowland, 2020b |
| Mean daily maximum temperature per month | tmax | °C | Met Office, 2022 |
| Mean daily minimum temperature per month | tmin | °C | Met Office, 2022 |
| Number of days with air frost per month | af | days | Met Office, 2022 |
| Total rainfall per month | rain | mm | Met Office, 2022 |

To the data frame containing information on the date of spawning, we added the data on maximum ammonium and nitrate ion concentrations recorded in each pond in each year whilst breeding adults were present. To the data frames containing information on the surface area of spawn, the percentage of spawn dead, and the date of hatching, we added data on mean ammonium and nitrate ion concentrations recorded in each pond in each year whilst spawn was present. To the data frame containing information on the date of leaving, we added data on the mean ammonium and nitrate ion concentrations recorded in each pond in each year whilst tadpoles were present.

To calculate land cover change, we made assumption that the *rate* of land cover change was constant between years, allowing us to estimate the area of each land cover class surrounding each ECN location in each year from 1994-2015 (Table 2). We identified the nearest Met Office station to each ECN location and used the climate data from these stations as estimates for the climate at the ECN locations. As congregation and spawning dates depend on winter climate (Carroll *et al*. 2009; Benard 2015), we calculated the average of each climatic variable at each ECN location in each year over the winter months (December-March). We calculated the average of each climatic variable at each ECN location in each year during the months whilst spawn was present (January-May) and added these values to the data frames containing information on the surface area of spawn, the percentage of spawn dead, and the date of hatching. Finally, we calculated the average of each climatic variable at each ECN location in each year during the months whilst tadpoles were present (February-September) and added these values to the data frame containing information on the date of leaving.

### Data Analysis

All data and code to reproduce these analyses is provided at https://github.com/xavharrison/FrogSpawn2023

As landscape-scale variables such as land cover, temperature and chemical use are often correlated, we used a Principal Components Analysis (PCA) on the anthropogenic stressor variables for each of the reproductive success and phenology data frames using the factoextra and FactoMineR packages (Sebastien Le 2008; Mundt 2020). We extracted Principal Component 1 (PC1) and Principal Component 2 (PC2) for each reproductive success and phenological variable to use as predictors in our modelling. We used rotation plots to identify the anthropogenic stressors explained by each Principal Component (Simko 2021), and so crucially the biological interpretation of the importance of each PC changes for each model.

We used Generalised Linear Models (GLMs) and General Linear Mixed Effects Models (GLMM) to investigate drivers in variation in date of congregation (n=88), spawning (n=23), and hatching (n=25), as well as the surface area of spawn (n=34), and the percentage of dead spawn (n=33). Variation in the size of the datasets is a result of ensuring all rows have complete information on all traits, including land use change and pollution. For these datasets, we fitted GLMMs with Gaussian errors, with PC1 and PC2 as the explanatory variables, and the ECN location and/or pond as a random intercept. We logit-transformed the percentage of dead spawn prior to model fiwng. These models also controlled for temporal autocorrelation in residuals. We used the ‘Leave One Out’ Information Criterion (LOO-IC, Vehtari *et al*. 2017) to identify the model(s). We performed ‘full model tests’, comparing the full model against a null model, to minimise bias in standard errors / credible intervals and control the Type I error rate (Forstmeier & Schielzeth 2011).

Finally, where we identified reproductive success and phenological variables to be associated with temperature (i.e. temperature loaded significantly into PC1 or PC2), we used bivariate mixed effect models to estimate the posterior correlation between temperature and phenological outcome variables. That is, variation in temperature arises due to latitudinal gradients as well as climate change, and so a higher standard of evidence is required to link shifts in phenology specifically to climate change rather than differences in site location. These bivariate models quantify correlation among the *residuals* of the two responses (e.g. mean maximum temperature and hatch date), whilst also accounting for fixed effects such as year/trends over time (see Houslay & Wilson 2017). Here we would predict that higher than average maximum temperatures (a positive residual) would be associated with earlier than average hatch dates (a negative residual). This manifests as a negative correlation in the residuals of the model, assesses as significant/important depending on whether the credible intervals cross zero. This approach is similar to ‘detrending’ residuals to identify causal effects (e.g. see Votier *et al*. 2008)).

## RESULTS

### Congrega1on Date

We detected a negative relationship between PC1 and congregation date, meaning that higher maximum temperatures, and higher proportions of urban and arable area were associated with earlier congregation to breed (Fig. 2A,B). There was no support for an effect of PC2 (credible intervals crossed zero). The full model was a superior fit to a null model (ΔLOO-IC 49.7). The model explained 81.5% of variation in congregation date [95% CI 77.2 = 84.3%]. A bivariate model estimating the strength of correlation between maximum temperature and congregation date whilst controlling for temporal autocorrelation supported these patterns, where higher than average temperatures are associated with earlier than average congregation to breed (mean correlation -0.52, 95% CI -0.01 --0.85; Fig 2C).

**Figure 2.**
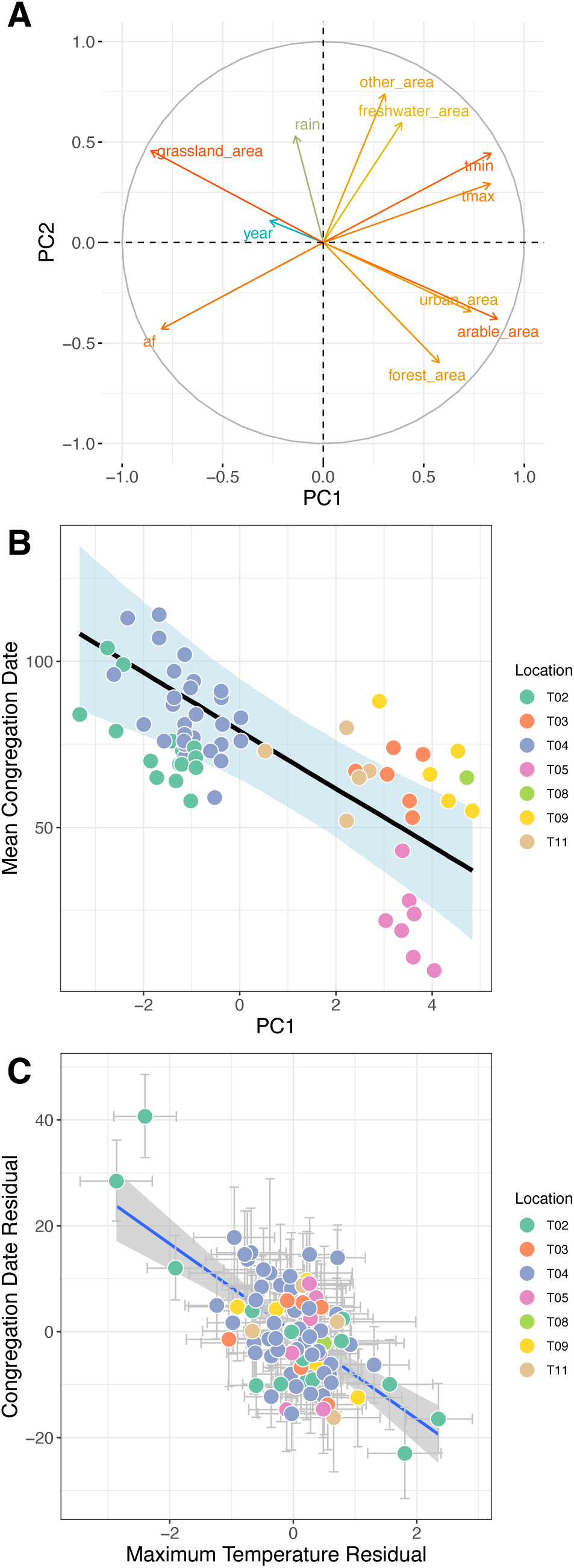
(A) Loadings of input variables on the first 2 axes of a Principal Component Analysis explaining variation in congregation date. (B) Significant negative relationship between PC1 and congregation date (C) Significant negative correlation between maximum temperature at site and congregation date.

### Spawn Date

As for congregation date, we uncovered a negative relationship between PC1 and spawn date. Higher maximum temperatures, and higher proportions of urban and arable area were associated with earlier spawning (Fig. 3A,B). There was no support for an effect of PC2 (credible intervals crossed zero). The full model was a superior fit to a null model (ΔLOO-IC 40.3). The model explained 87% of variation in spawn date [95% CI 77.3 = 92%]. A bivariate model estimating the strength of correlation between maximum temperature and spawn date whilst controlling for temporal autocorrelation supported these patterns (mean correlation -0.67, 95% CI -0.34 --0.87; Fig 3C).

**Figure 3.**
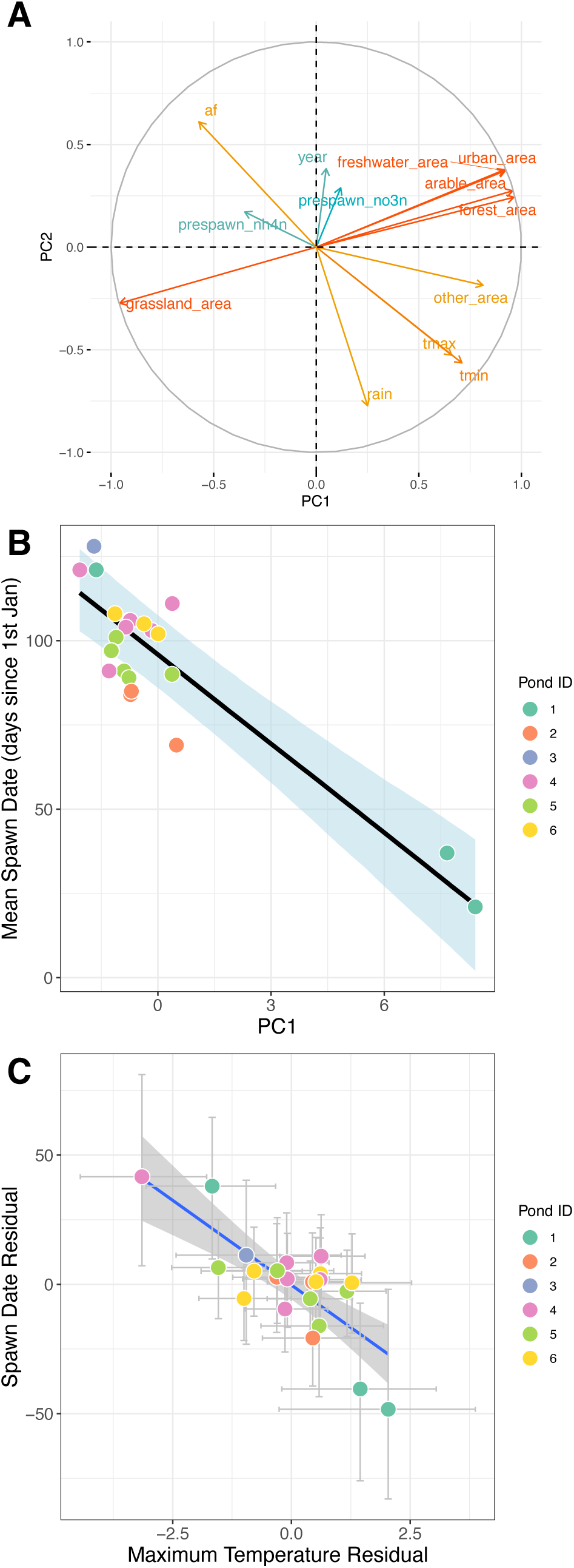
(A) Loadings of input variables on the first 2 axes of a Principal Component Analysis. (B) Significant negative relationship between PC1 and spawn date (C) Significant negative correlation between maximum temperature at site and spawn date.

### Hatch Date

For hatch date, we detected significant effects of PC2, where increases in freshwater area and higher *minimum* temperatures were associated with earlier hatching, and an increased in the number of Air Frost days was associated with later hatching (Fig. 4A; Fig. S1). A model containing effects of PC1 and PC2 was marginally superior fit to an intercept only model (ΔLOO-IC = 6.8; Fig. 4B). There was no support for an effect of PC1 (credible intervals crossed zero).

**Figure 4.**
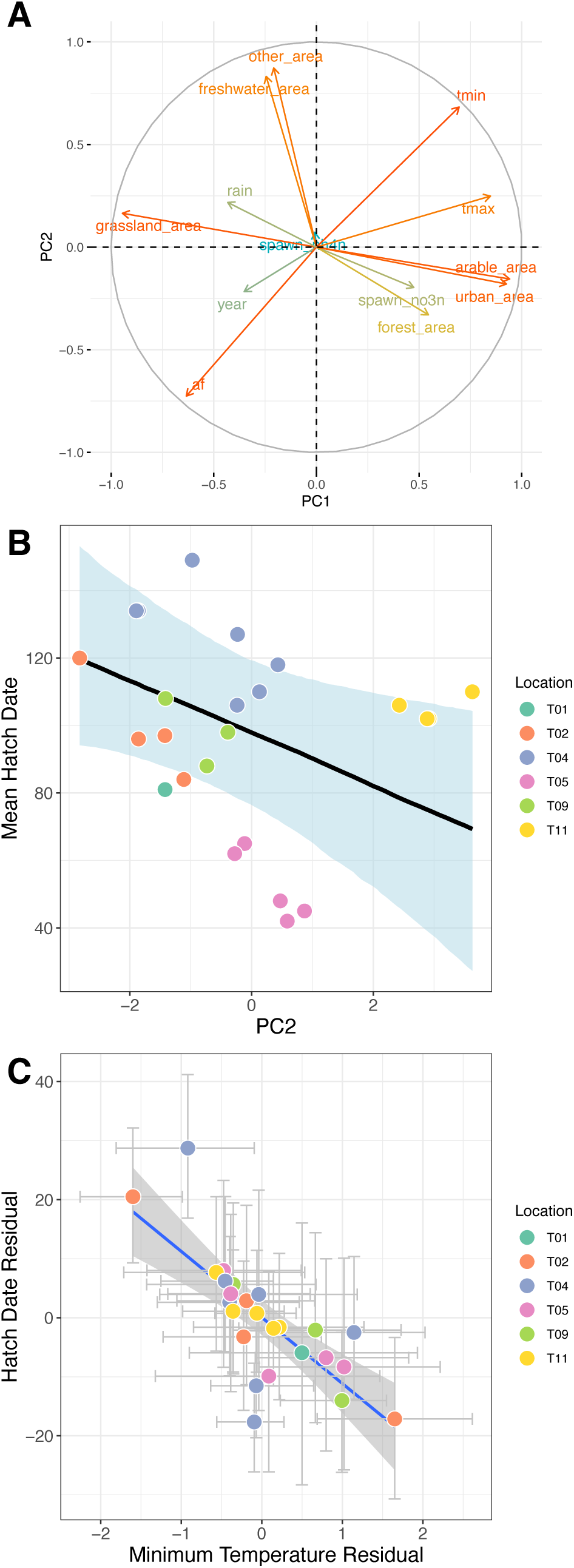
(A) Loadings of input variables on the first 2 axes of a Principal Component Analysis. (B) Significant negative relationship between PC2 and hatch date (C) Significant negative correlation between maximum temperature at a site and hatch date.

The model explained 89% of variation in hatch date [95% CI 77.3 = 92%]. A bivariate model estimating the strength of correlation between minimum temperature and hatch date whilst controlling for temporal autocorrelation supported these patterns (mean residual correlation -0.65, 95% CI -0.27 --0.88; Fig 4C). A bivariate model with maximum temperature as a predictor returned similar results.

### Reproduc1ve Success

We detected a negative association between PC1 and the percentage of dead spawn observed (Fig 5A,B). PC1 represents a metric of the area of arable, freshwater, grassland, and built-up land cover, and the mean maximum daily temperatures spawn was exposed to (Fig 5A). Higher cumulative days of air frost and higher grassland area were associated increased percentages dead spawn (Fig. S2), whilst increases in arable area, built-up area, and maximum temperature led to reductions in the percentage of dead spawn (Fig 5B; Fig S2). This model explained 45% of the variance in percentage of dead spawn (95% CI 22.5 – 60.8%]. Unlike previous traits, there was no support for a relationship between maximum temperature and % dead spawn (credible intervals crossed zero, though the relationship was negative (mean correlation -0.28, 95% CI -0.58 – 0.09). Finally, there was no statistical support for an effect of any of variable on the maximum surface area of spawn observed.

**Figure 5.**
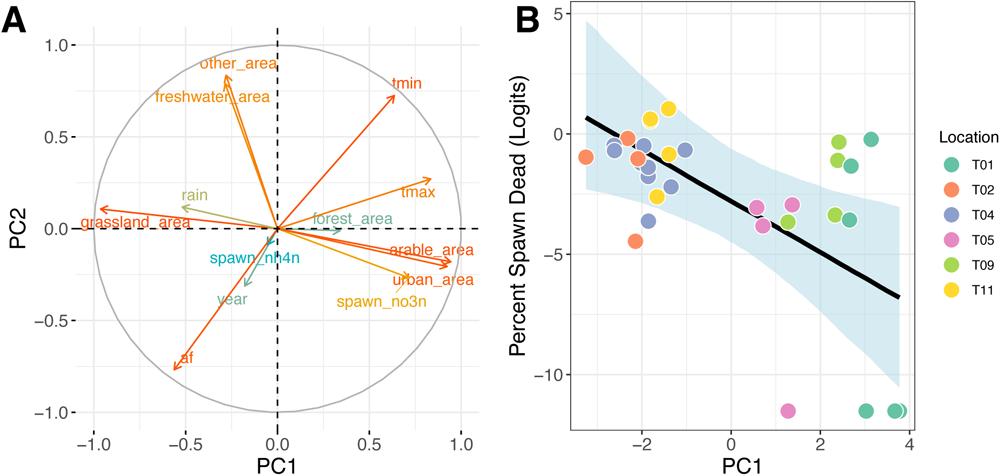
(A) Loadings of input variables on the first 2 axes of a Principal Component Analysis. (B) Significant negative relationship between PC1 and the percentage of spawn dead.

## DISCUSSION

Here we used long-term data on the breeding behaviour of UK *Rana temporaria* to identify associations with land use change, pollution, and climate. We found that higher maximum temperatures were associated with earlier congregation & spawning, whilst higher minimum temperatures were linked to earlier hatching. No predictors could be linked to the surface area of spawn present at a site, but higher numbers of frost days were associated with higher proportions of dead spawn. All reproductive traits we measured could also be linked to traits of land use change and pollution (via Principal Component Analysis), which themselves correlated with temperature variation. This study highlights the important role of winter temperature in driving variation in amphibian breeding phenology, but also the complexity of disentangling the relative significance of multiple correlated environmental variables in wild phenology studies.

### Phenology, Land Use Change and Temperature

This work is consistent with previous studies linking temperature shifts to alteration in amphibian breeding phenology (Reading 2007; Blaustein *et al*. 2010; Benard 2015). Consistent with our predictions, we found that warmer average temperatures correlated with advancements in the timings of adult congregation to breed, spawning, and hatching. Crucially, our modelling approach allows us to disentangle the relative effects of latitudinal variation in breeding phenology and *within loca1on* variation caused by local shifts in climate, such as mean winter temperature. These bivariate models revealed that within sites, warmer than average winter temperatures were associated with earlier than average metrics of breeding phenology. Taken together these data suggest that *R. temporaria* breeding phenology shows a degree of plasticity, and responds to local variation in temperature among years by shifting their timing of congregation, spawning and hatching. Similar patterns have recently been observed in two newt species (Hubáček & Gvoždík 2023), suggesting this many amphibian species are capable of exhibiting such individual plasticity.

Warmer winters have been shown to advance phenology in congeners like wood frogs (*Rana sylva1ca*), but also delay larval development time (Benard 2015). Crucially earlier breeding does not compensate for this developmental delay. In common toads *Bufo bufo* in the United Kingdom, warmer than average temperatures correlated with decreased body condition and survival (Reading 2007). Recent work on wood frogs showed that phenological shifts can expose individuals to colder temperatures and resulted in lower tolerance of offspring to pollutants like NaCl (Buss et al 2021). Collectively these data suggest that in some species, warming can induce cryptic cost of breeding plasticity in multiple life stages that are not immediately apparent if looking at phenological variables alone (see Blaustein et al 2010).

We also uncovered associations between land use change and breeding phenology. Increased arable and built-up land cover is associated with earlier hatching, whilst grassland is associated with later hatching. Arable and built-up areas tend to be warmer than the surrounding natural habitats due to unvegetated ground, whilst grassland areas, with higher vegetation levels, are cooler (Lembrechts *et al*. 2019; Schmidt *et al*. 2019). The warming effects of arable and built-up areas can even increase the temperature of the wider landscape, leading to earlier breeding phenology in the areas surrounding human-dominated land (Tian *et al*. 2020). These effects could be reduced by increasing vegetated areas in these land cover types, thus providing cooler microclimates (Greenwood *et al*. 2016).

### Land Use Change, Temperature and Spawn Mortality

We found no clear association between mean winter maximum or minimum temperatures and spawn mortality, when temperature was used as the sole predictor in models. Instead, we found that the percentage of dead spawn was linked to a composite measure of frost days, rainfall, land use change, and temperature. Increases in the proportion of arable and urban areas, nitrate levels, and increased mean maximum temperature, were associated with higher spawn survival. Conversely, increased numbers of frost days, rainfall and grassland areas were associated with decreasing reproductive success.

These associations did not align with our predictions, where we expected increased farmland (and associated pollution such as nitrates), and increased urbanisation to be detrimental rather than associated with higher spawn survival. However, though human-driven land use change is often associated with lower reproductive success, some studies have shown that they can support declining populations. For example, the average reproductive success of multiple amphibian species in America breeding in arable ponds was no different to in natural wetlands (Knutson *et al*. 2004). Though *some* species did respond negatively to arable land use (Knutson *et al*. 2004), *R. temporaria* have been found to use arable ponds more than other amphibians (Hartel *et al*. 2011). Therefore, arable land cover may be able to support robust populations of *R. temporaria*, leading to higher spawn survival in the ECN sampling locations. Additionally, arable ditches have been shown to confer landscape connectivity for amphibians, reptiles, and mammals (Maisonneuve & Rioux 2001; Jobin *et al*. 2004). Maes *et al*. (2008) even found that ditches in agricultural environmental schemes supported similar *Rana esculenta* abundances as nature reserves, demonstrating how these features of arable land cover can serve as important avenues for dispersal between breeding sites. This is crucial in preventing population isolation, perhaps explaining why we found a negative relationship between the arable area and the maximum percentage of spawn that died (Allentoft & O’Brien 2010).

Built-up areas have also been shown to increase the fitness of some wildlife populations, with some threatened birds (Kettel *et al*. 2019) and amphibians (Iglesias-Carrasco *et al*. 2017) having greater reproductive success ad body condition respectively in built-up areas than in the countryside. There are multiple theories for why this could be. Saenz *et al*. (2015) suggest that the occurrence of chytridiomycosis, an amphibian disease that can reduce fitness, could be lower in urban areas. However, chytridiomycosis is not common in British amphibians (Garner *et al*. 2005) so this is unlikely to explain our findings. Hall and Warner (2017) suggest that the high densities of prey insects in urban areas or lower predation pressures that allow adults to spend more time hunting, could increase fitness. Further evidence for this is from Germany, where *R. temporaria* adults were found to be bigger in urban greenspace than in the surrounding countryside (Niemeier *et al*. 2020). Larger adult body size is likely to increase offspring survival (Hall & Warner 2017), providing an explanation as to why built-up areas are associated with lower *R. temporaria* spawn mortality.

Many of the studies that demonstrate the value of human-dominated land cover types also acknowledge the need for management. In arable areas, vegetation complexity in ditches should be encouraged (Maisonneuve & Rioux 2001) to allow ditches to decrease *R. temporaria* population isolation. Similarly, in built-up areas, it is important to reduce barriers to movement by connecting urban greenspace (Mazgajska & Mazgajski 2020; Niemeier *et al*. 2020). These suggestions highlight that local management for *R. temporaria* could be more important that the broad-scale land cover type in determining this amphibian’s reproductive success.

Landscape management practices could also explain why grassland is negatively associated with *R. temporaria* reproductive success. In the UK, only 2% of grassland is classed as diverse (Bullock *et al*. 2011), due to widespread “improvement” (Vickery 1999) and livestock grazing (Fuller 1987; Bullock *et al*. 2011). However, grazing can be detrimental to amphibians. Livestock can cause high levels of wetland bank erosion (Trimble 1994), leading to increased sediment deposition. When investigating the impact of cattle on *Bufo achalensis*, a toad species endemic to Argentina, Jofré *et al*. (2007) found that increased sediment levels reduced algal growth, a key food source for larval amphibians, leading to higher mortality in the *B. achalensis* larvae. This could result in increased isolation between *R. temporaria* populations, potentially leading to the increased percentage of spawn death observed in ECN locations surrounded by grassland. High densities of livestock can also directly reduce amphibian spawn reproductive success through disruption and trampling (Knutson *et al*. 2004).

The negative relationship between freshwater area and reproductive success seems counterintuitive, however, predator presence could explain this relationship. The land cover data used in this study had a resolution of 25x25m, so only large water bodies were present in the data (Rowland 2020a, b). Previous studies have found that these are often avoided by amphibians are they are more likely to contain a high density of predators (Pearman 1995). Knutson *et al*. (2004) even found that the presence of predatory fish was one of the most important factors in determining the reproductive success of an American amphibian community. Therefore, the presence of large areas of freshwater surrounding the ECN sampling locations may increase the isolation of ECN breeding populations, resulting in reduced spawn survival and higher spawn mortality.

Our modelling could not conclusively implicate temperature regime changes as a potential driver of difference in amphibian reproductive success, though maximum temperature and number of frost days were part of the composite measure (PC1) associated with spawn survival. Although climate change is leading to warmer temperatures, it is also causing extreme weather conditions (Huber & Gulledge 2011). One example is the occurrence of spring cold-snaps, a phenomenon that has been shown to have detrimental effects on a wide range of organisms (Augspurger 2013; Benard 2015; Turner & Maclean 2022). Benard (2015) observed an increase in *Rana sylva1ca* larvae being exposed to cold-snaps from 2006 to 2012, leading to altered development. Freezing is known to kill *R. temporaria* (Pasanen & Karhapää 1997), making it likely that cold-snaps could lead to high spawn mortality. Warmer springs and fewer frost days likely explain the lower proportions of dead spawn observed in this dataset under these conditions.

### Conclusions and Future Work

Here we have demonstrated associations between climate, land use change and parameters of amphibian breeding success and phenology at the landscape scale. Our results are consistent with investigations at the scale of individual ponds that uncovered similar relationships between temperature and the timing of amphibian breeding and larval development (Benard 2015). Future work on *R. temporaria* in the UK should prioritise pond-scale approaches to these questions, which will permit measurement of microhabitats experienced by breeding adults. Microhabitat temperature measurements could shed further light on the frequency and consequences of freezing temperature (i.e. lower winter minima) on *R. temporaria*, as well as their effect on larval development and survival. Similarly, fine-scale land-use data could aid in identifying the habitats most beneficial for successful *R. temporaria* breeding, allowing the efficacy of management practices to be optimised. The need for fine-scale data is particularly important due to the small size and limited dispersal ability of *R. temporaria* (Kovar *et al*., 2009), and thus may also be applicable to other such species. For the ongoing survival of *R. temporaria* it is vital to reduce the harm of extreme weather events due to climate change, which may be achieved through the management of microclimates. Afforestation and re-flooding drained wetlands, for example, could help maintain stable and favourable microclimatic envelopes. These practices could also aid *R. temporaria* by increasing vegetation complexity and providing additional breeding locations and their connectivity, both of which could improve *R. temporaria* survival.

## Supporting information

Supplementary Graphs

