## Supplementary Graphs for "Spring temperature and land use change are associated with *Rana temporaria* reproductive success and phenology"

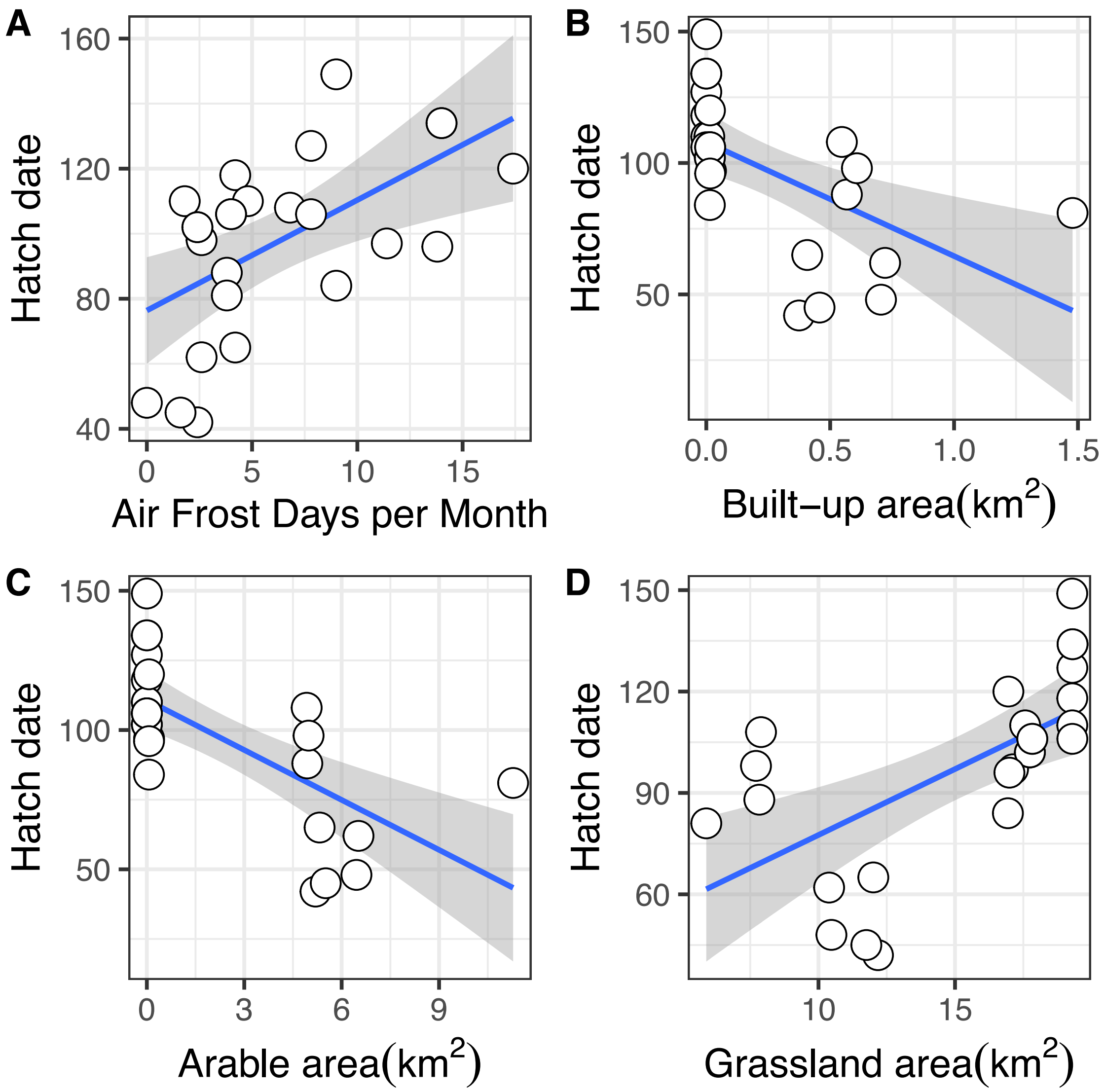


**Figure S1.** Factors associated with variation in the hatch date. More air frost days (A) resulted in later hatching. Higher built up area (B) and arable area (C) were associated with earlier hatching, whereas increased grassland area (C) was linked to later hatching.

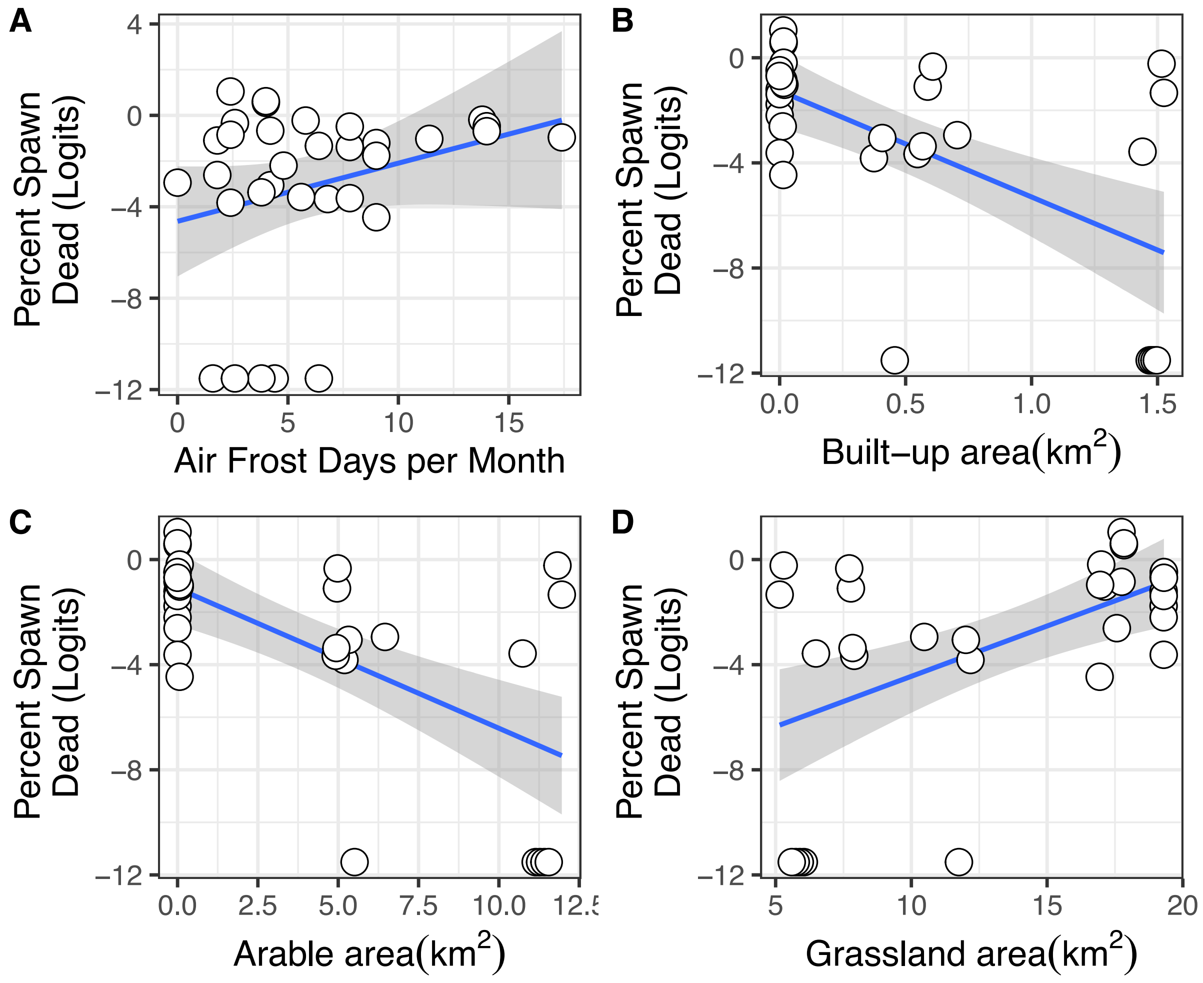


**Figure S2.** Factors associated with variation in the percentage of dead spawn. More air frost days (A) resulted in higher death. Higher built up area (B) and arable area (C) were associated with higher spawn survival, whereas increased grassland area (D) was linked to larger proportions of dead spawn.
